# Biological age acceleration is more prevalent in men than in women at time of spontaneous intracerebral hemorrhage

**DOI:** 10.1101/2025.10.01.679916

**Authors:** Cristina Gallego-Fabrega, Joan Jiménez-Balado, Natalia Cullell, Jara Cárcel-Márquez, Elena Muiño, Laia Llucià-Carol, Jesús M. Martin-Campos, Laia Mariné, Paula. Villatoro-Gonzalez, Paula Boldo, Ana Aguilera-Simon, Anna. Ramos-Pachon, Pol. Camps-Renom, Luis. Prats-Sanchez, Jaume Roquer, Joan Martí-Fabregas, Jordi Jiménez-Conde, Israel Fernández-Cadenas

## Abstract

**Objective:** Spontaneous intracerebral hemorrhage (ICH) occurs later in life in women compared to men. Although previous studies have demonstrated epigenetic age acceleration (EAA) differences in ischemic stroke (IS) patients - a measure of an individual’s biological aging -it remains unknown whether this sex dimorphism also applies to ICH patients.

**Approach and Results:** We joined two ICH cohorts (N= 200, 45% women: 76.8±13 years; 55% men: 67.9±14 years). DNAm levels were obtained from whole blood samples using Illumina EPIC array. We evaluated three age-predictor clocks (Horvath, Hannum and Zang-BLUP) and one health-status clock (Levine) and their respective EAA metrics including extrinsic EAA (EEAA) and intrinsic EAA (IEAA) measures. We compared aging measures between women and men, then performed an ICH-subtype stratification, a specificity analysis evaluating sex differences in non-stroke samples (N= 350, 54% women 60.8±8 years, men: 60.8±10 years) and replication of previous result in a new IS cohort (N= 657, 43% women 73.6±12 years, men: 70.1±11 years).

Women at time of ICH present lower EAA values than men (Horvath-EAA, p-value=1×10^-04^; Hannum-EAA, p-value=8.1×10^-06^; BLUP-EAA, p-value=4.4×10^-04^) as well as lower extrinsic EAA values (Horvath-IEAA, p-value=3.1×10^-03^; Hannum-IEAA, p-value=6.5×10^-04^). These differences seemed to be driven by differences in deep-ICH patients. Non-ICH females have lower acceleration values (Hannum-EAA: -10,37) than non-ICH men (Hannum-EAA: -8,01), but the differences are smaller than in ICH cases (ICH-women Hannum-EAA: -9,26, ICH-men Hannum-EAA: -2,6). This pattern was consistent in the IS cohort, where women were chronologically older than men but had similar biological age and significantly lower epigenetic age acceleration across multiple measures (Horvath-EAA, p = 2.6×10^−3^; Hannum-EAA, p = 7.1×10^−3^; Horvath-IEAA, p = 1.6×10^−2^; Hannum-IEAA, p = 4.5×10^−2^).

**Conclusion:** This study shows that biological age difference between women and men are not exclusive of ischemic stroke but also observed in ICH.

## INTRODUCTION

Intracerebral hemorrhage (ICH) is a type of stroke that occurs when a blood vessel ruptures and bleeds into the brain parenchyma. It accounts for up to 27,9% of all strokes worldwide[1] and is associated with high mortality and disability[2]. Although absolute numbers of strokes cases have increased over the last 30 years, the age-standardized rates have decreased[1].

There is evidence that men and women who experience ICH differ in characteristics, risk factors, severity, outcome, and treatments[3]. Women are, on average, older than men at time of ICH[4, 5], with worse previous neurological status, and higher proportion of deep ICH compared to men. Men, however, have more risk factors, such as a more frequent presence of high blood pressure, obesity, smoking habit, as well as early hematoma expansion and higher 90 day and one year mortality rates[6].

Age at ischemic stroke onset is generally younger in men than in women[1]. A previous study by our group reported that, despite this difference in chronological age, men and women have the same biological age at the time of ischemic stroke (IS)[7]. Our study revealed that women exhibited negative age acceleration (or deceleration), while men showed the opposite trend[7].

Biological age describes how old a person’s cells or tissues are based on physiological evidence. One of the most common biological age estimators is obtained through DNA methylation (DNAm) data[8–10]. Several authors have been able to identify DNAm sites highly associated with aging rate or health status in what are called Epigenetic Clocks[11, 12]. Epigenetic clocks are reported to be better predictors of IS risk, outcome and mortality, than chronological age[13–15], as well as a host of other age-related diseases[16, 17]. DNAm measures, independent of chronological age, known as epigenetic age acceleration (EAA)[12, 18, 19], are obtained after regressing biological age on chronological age representing the rate of aging.

Unlike DNA mutations, DNAm changes are reversible making them an attractive target for developing therapies aimed to slow down the aging process[8] and thus the risk of age-related diseases like ICH. It has been shown that the use of medications commonly used to treat chronic diseases like hypertension and hyperlipidemia can reduce measures of age acceleration [20], as well as adherence to a Mediterranean diet[21].

Based on the described sex-dimorphisms in ICH patients, here we aim to interrogate whether age acceleration is an underlaying mechanism driving such dimorphism and to examine whether those differences are dependent on hemorrhage localization.

## MATHERIAL AND METHODS

### 2.1. Participants

#### Data Availability

Data supporting these findings is available from the corresponding authors upon reasonable request.

#### Study Sample

A total of 200 ICH patients from two Spanish hospitals (Hospital del Mar, Barcelona and Hospital de la Santa Creu I Sant Pau, Barcelona), where analyzed. We selected first ever ICH cases >18 years-of-age with data on age, sex, smoking status, diabetes, dyslipidemia, hypertension and hemorrhage location, and available biological material to perform an epigenomic study (Table 1). Detailed information about hemorrhage location can be found in the supplemental material (Supplemental Table S1).

**Table 1:**
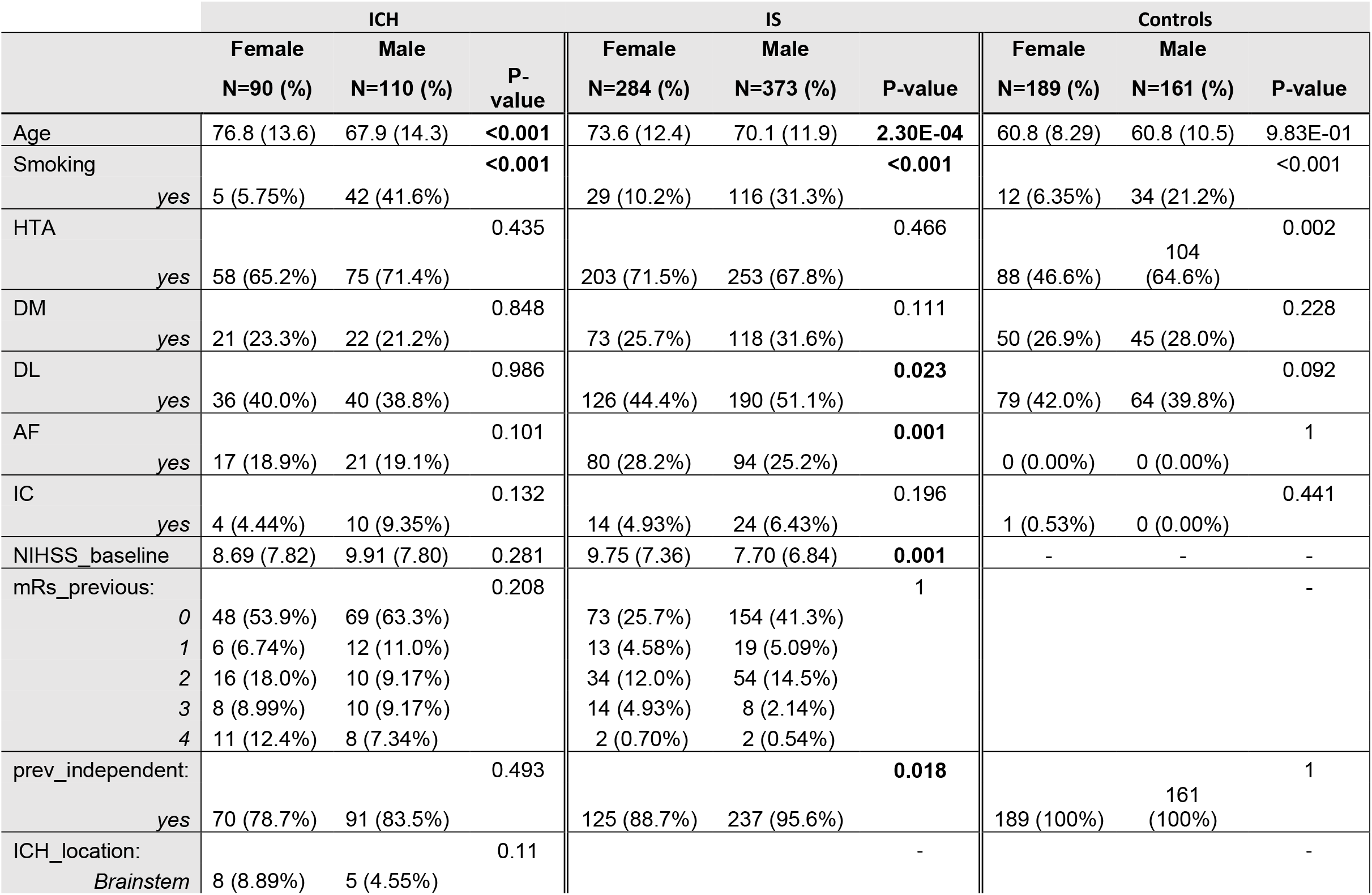
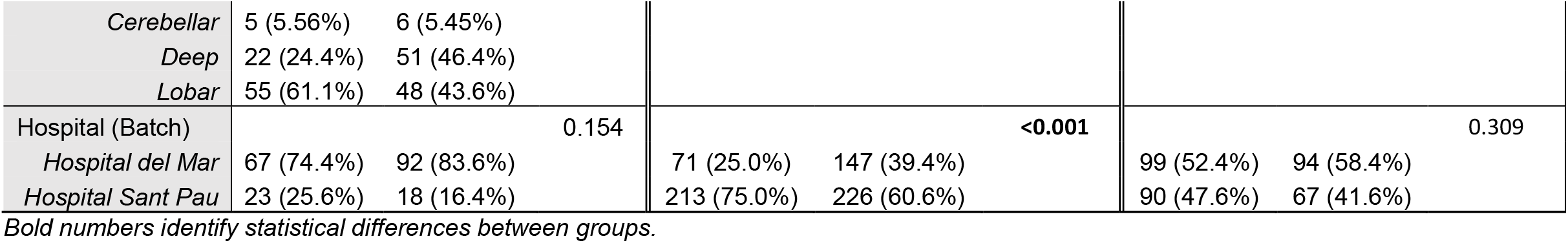
Demographic characteristics or the study cohort.

As a specificity test and to replicate previous results, we also analyzed 350 non-stroke controls and 657 new IS samples from the same hospitals, respectively. Inclusion criteria for IS cases were >18 years-of-age with data on age, sex, smoking status, diabetes, dyslipidemia, hypertension and stroke subtype (Table 1). For control samples were >18 years-of-age without previous cerebrovascular events and data on age, sex, smoking status, diabetes, dyslipidemia and hypertension (Table 1).

The study was carried out in accordance with the Declaration of Helsinki and approved by the respective hospitals’ research ethics committees: institutional review board at the Hospital del Mar (CPMP/ICH/135/95) and institutional review board at the Hospital de la Santa Creu i Sant Pau (COLECCION 03/2016). Written informed consents were obtained from all participants.

#### Sample Collection

In Hospital del Mar ICH and IS blood samples were collected within the first 24 hours of stroke onset, DNA was extracted using manual salt precipitation in the Banco Nacional de ADN (Instituto de Salud Carlos III) and stored at –80ºC until utilization. ICH samples from Hospital Sant Pau were collected during hospitalization within 14 days of stroke onset and IS samples within the first 24 hours of stroke onset. DNA was extracted using the Gentra Pure gene Blood Kid (Qiagen, Hilden, Germany) following the manufacturer’s instructions and stored at −80ºC until required.

### 2.2. Methylation Study

#### Epigenome Wide Methylation Assay

Genome-wide DNAm was evaluated using the Infinium MethylationEPIC BeadChip (EPIC) (Illumina Inc, San Diego, CA), which analyzes more than 850,000 CpG sites (CpGs)[22]. DNA methylation values (β-values) were estimated using the GenomeStudio Software (Illumina).

The success of bisulfite conversion, array hybridization and sample preparation were evaluated through comprehensive quality control (QC) metrics, using *ChAMP* (v.2.26.0) R package[23], following established protocols[24, 25]. In short, samples presenting sex-mismatch between genotypically obtain information and medical record were removed. Follow by removal of probes with unsuccessful hybridization (detection p-value>0.05) and lacking representation (<3 beads/probe). To address issues related to poor bisulfite conversion, samples were excluded if more than 1% of probes exhibited a detection p-value>0.01.

### 2.3 DNAm-based Biological Age Calculation

Biological age estimators and age accelerations were calculated using the *methylclock*^19^ (v.1.8) R package[19]. We selected four complementary clocks to capture different dimensions of biological aging and to improve robustness and generalizability of findings. These included: Horvath’s multi-tissue epigenetic clock[26] (Horvath-BioAge), Hannum et al.’s blood-based clock[27] (Hannum-BioAge), and Zhang et al.’s Best Linear Unbiased Prediction model[28] (BLUP-BioAge), which are widely used DNA methylation-based predictors of chronological age. We also included Levine’s PhenoAge[29] (Levine-BioAge), which is designed to predict morbidity and mortality risk, making it a health-span and lifespan indicator.

For each clock, we also obtained epigenetic age acceleration (EAA) measures. Regular EAA (Horvath-EAA, Hannum-EAA, BLUP-EAA, Levine-EAA) reflects the residuals from regressing DNAm-age on chronological age, and indicates whether an individual’s biological age is higher or lower than expected. Intrinsic EAA (IEAA) (Horvath-IEAA, Hannum-IEAA, BLUP-IEAA, Levine-IEAA) adjusts for in-silico derived blood cell-type composition and isolates “pure” epigenetic aging effects, largely independent of immune cell changes. Extrinsic EAA (EEAA) (Hannum-EEAA, BLUP-EEAA, Levine-EEAA) incorporates age-related changes in blood cell types and is thought to better capture immune system aging and systemic inflammation. As Horvath’s clock was trained in samples from multiple tissues, is not specific for blood samples and therefore EEAA won’t be estimated Including these variants allows us to differentiate between cell-intrinsic epigenetic aging (IEAA) and immune-related aging processes (EEAA), which may have distinct biological and clinical implications.

### 2.4 Statistical Analyses

The R statistical computing environment[30] (v.4.2.1) was used to perform QC and preprocessing, as well as all statistical analyses and figures.

Associations of known cerebrovascular risk factors and ICH location with sex were assessed using t-test for quantitative dependent variables and chi-square for categorial ones (Table 1). Based on the univariate analysis results we constructed two multivariate generalized linear models, to adjust potential covariates. Model 1 (Model.1) included smoking status, and Model 2 (Model.2) was Model.1 plus ICH location. Additionally, in the case of IS samples we created the third model (Model.3) including all potential confounding factors identified in the univariate analysis (Table 1) (smoking, dyslipidemia, atrial fibrillation, baseline NIHSS and pre-stroke functional independence); similarly for control samples we created model four (Model.4) which included smoking and hypertension.

The fitness of the obtained biological clock’s measures was evaluated using Pearson correlation test, between chronological and biological age. To account for multiple testing, we applied a Bonferroni correction based on the four independent clocks evaluated (*p* < 0.05/4 = 0.0125). Acceleration measures are not independent off each other thus were not accounted for in the multiple comparison threshold.

#### Stratification analysis

To interrogate the biological age differences between men and women based on ICH location, we perform a stratification analysis by studying independently deep, lobar, brainstem and cerebellar ICH cases. For which we use the same strategy as the main analysis; univariate analysis to identify potential confounders of each sub-cohort, and a multivariate linear model accounting for smoking habit (Model.2) for consistency with the main analysis (Supplemental material Table S1).

### 2.6. Reporting Checklist for Cohort Study

STROBE (Strengthening the Reporting of Observational Studies in Epidemiology)[31] and SAGER (Sex and Gender Equity in Research)[32] reporting guidelines can be found in the supplemental material.

## RESULTS

We evaluated three biological age and one health span predictors in ICH patients, all of them were highly correlated with chronological age (Supplemental material Figure S1) (Horvath-BioAge: ρ=0.79, p-value = 5.5×10^-43^; Hannum-BioAge: ρ=0.84, p-value = 5×10^-54^; BLUP-BioAge: ρ=0.77, p-value = 5.5×10^-40^; Levine-BioAge: ρ=0.91, p-values = 4.1×10^-78^). Although women were chronologically older than men at time of ICH onset, no differences between sexes were observed in biological age in none of the measures in our regression models prior or after multiple-testing comparisons (Tabe 2) (Model.2: Horvath-BioAge, p-value = 0.35; Hannum-BioAge, p-value = 0.4; BLUP-BioAge: p-value = 0.77; Levine-BioAge: p-value = 9.6). Extended results for the univariate and Model.1 analysis can be found in Supplemental material Table S2.

### Sex Dichotomy in Age Acceleration in ICH Patients

When assessing the basic EAA measures, we observed that women present lower acceleration values than men for the three DNAm clocks and the health span predictor (Figure 1, Table 2), although differences were less pronounced in the latter (Horvath-EAA, bonf.p-value = 1×10^-04^; Hannum-EAA, bonf.p-value = 8.1×10^-06^; BLUP-EAA, bonf.p-value = 4.4×10^-04^; Levine-EAA, bonf.p-value = 4.7×10^-01^). The same pattern was identified for IEAA and EEAA (Table 2, Supplemental material Figures S2 and S3). Statistically significant IEAA differences between women and men were exclusively for the DNAm clocks (Horvath-IEAA, bonf.p-value = 3.1×10^-03^; Hannum-IEAA, bonf.p-value = 6.5×10^-04^; BLUP-IEAA, bonf.p-value = 1.4×10^-02^; Levine-IEAA, bonf.p-value = 0.23); and EEAA differences were present only for Hannum measures (; Hannum-EEAA, bonf.p-value = 2.3×10^-03^; BLUP-EEAA, bonf.p-value = 0.1; Levine-EEAA, bonf.p-value = 1).

**Table 2:**
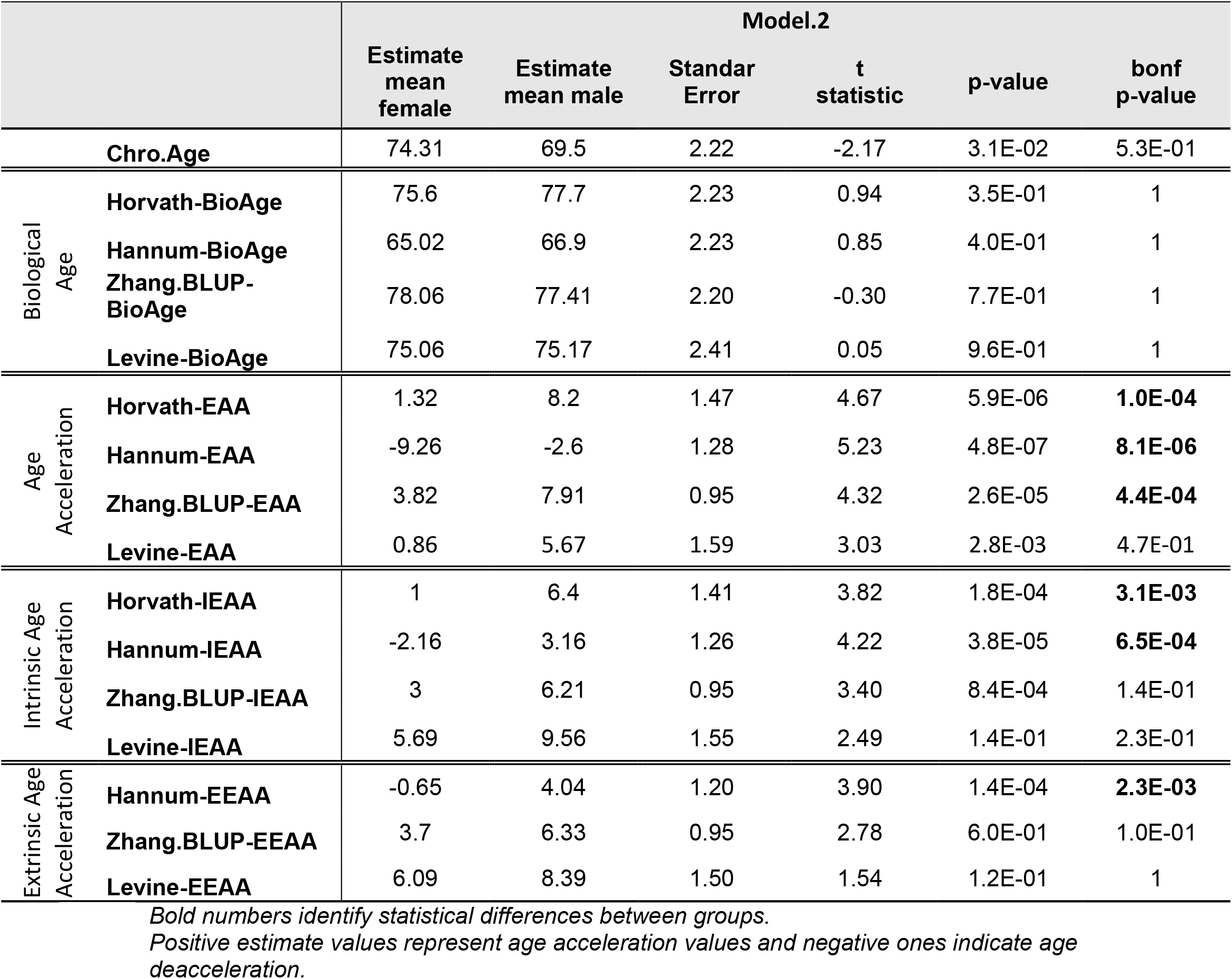
Differences in biological ages and acceleration measures between women and men, accounting for smoking habit and ICH location (Model.2).

**Figure 1:**
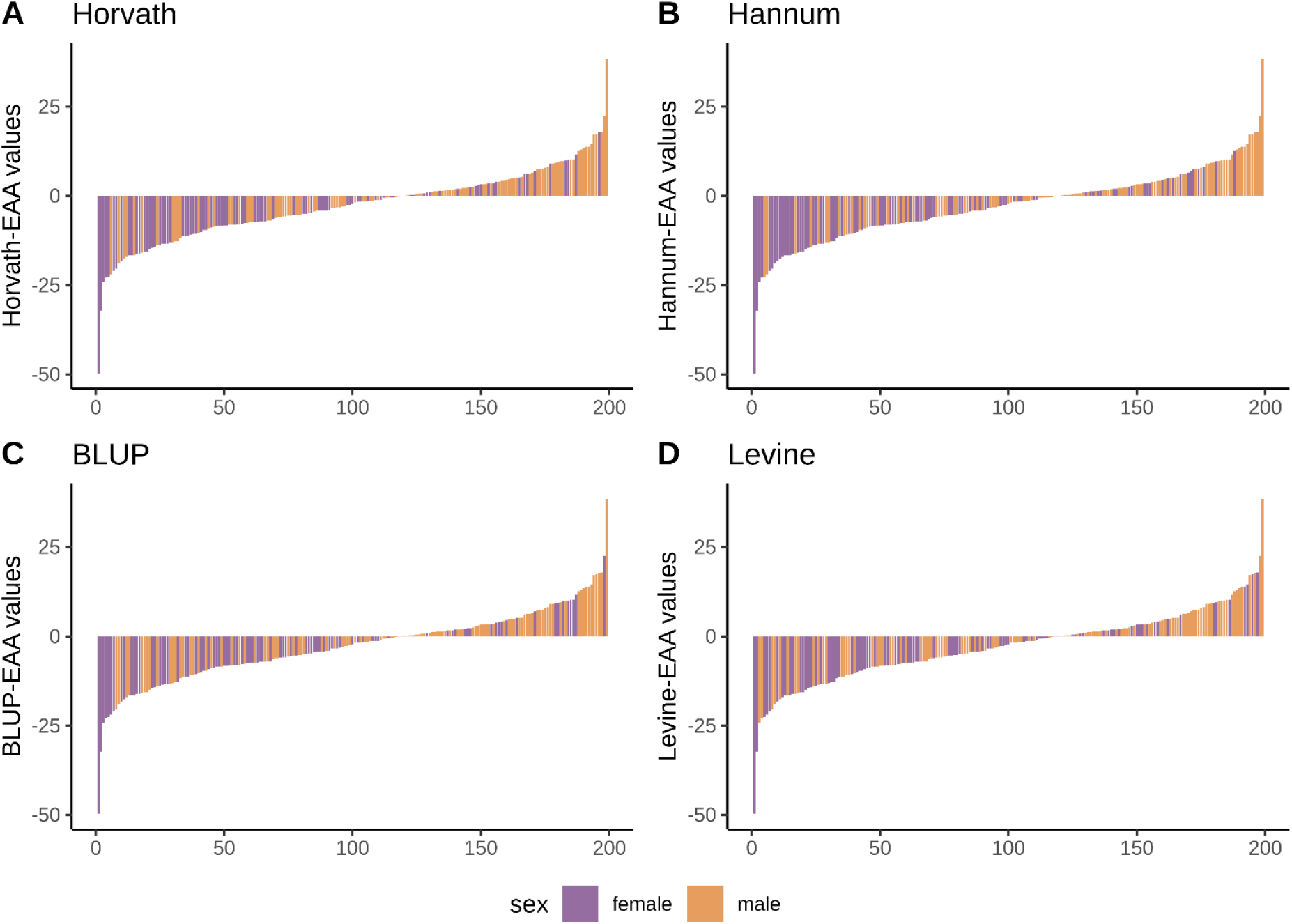
Distribution of EAA values between women and men across different DNAm Clocks. Negative values point out samples with negative acceleration, or younger than their chronological age. Positive values point out samples with positive acceleration or older than their chronological age. Each panel shows a different Epigenetic Clock (A: Horvath, B: Hannum, C: BLUP, D: Levine). Sample’s sex is differentiated by color, females: purple; males: orange. Each bar along X axes indicates a sample, Y exes indicates samples’ acceleration values. Ther is a higher rate of accelerate age values in man and a higher rate of deaccelerate age values in women *EAA: Epigenetic Age Acceleration*

A complete table with results from the univariate, Model.1 and Model.2 analysis can be found in Supplemental material Table S2.

### Sex Dichotomy in Age Acceleration stratified by ICH location

As per the sensitivity analysis, after the stratified analysis we observed the same patterns than in the full cohort, in which women tend to have lower acceleration values than men, although significant differences were only observed for the deep ICH subtype (Table 3), Horvath-EAA, bonf.p-value = 1.1×10^-03^; Hannum-EAA, bonf.p-value = 6.6×10^-03^; BLUP-EAA, bonf.p-value = 4.8×10^-03^). Additionally, we did a Hospital based stratification analysis in which the same trend was observed, which women show lower acceleration values than men, despite not reaching statistically significant differences (Supplemental material Table S3).

**Table 3:** Differences in biological ages and acceleration measures between women and men stratified by ICH location, accounting for smoking habit.

|  |  | Deep (Model.1) |  |  |  |  |  | Lobar (Model.1) |  |  |  |  |  | Brainsteam (Model.1) |  |  |
| --- | --- | --- | --- | --- | --- | --- | --- | --- | --- | --- | --- | --- | --- | --- | --- | --- |
|  |  | Estimate<br>mean<br>female | Estimate<br>mean<br>male | Standar<br>Error | t<br>statistic | p-value | bonf<br>p-value | Estimate<br>mean<br>female | Estimate<br>mean<br>male | Standar<br>Error | t<br>statistic | p-value | bonf<br>p-value | Estimate<br>mean<br>female | Estimate<br>mean<br>male | Standar<br>Error |
| Chro.Age |  | 75.82 | 67.19 | 4.07 | -2.12 | 3.80E-02 | 6.41E-01 | 74.94 | 71.18 | 2.73 | -1.38 | 1.70E-01 | 1 | 43.94 | 27.35 | 8.12 |
| Biological<br>Age | Horvath-BioAge | 61.38 | 65.54 | 3.93 | 1.06 | 2.90E-01 | 1 | 72.91 | 71.22 | 2.81 | -0.60 | 5.50E-01 | 1 | 66.68 | 55.30 | 8.07 |
|  | Hannum-BioAge | 59.03 | 60.23 | 4.01 | 0.30 | 7.60E-01 | 1 | 65.54 | 66.13 | 2.90 | 0.20 | 8.40E-01 | 1 | 41.08 | 31.89 | 6.20 |
|  | Zhang.BLUP-BioAge | 68.87 | 69.55 | 4.16 | 0.16 | 8.70E-01 | 1 | 77.82 | 75.03 | 2.71 | -1.03 | 3.10E-01 | 1 | 47.64 | 34.83 | 7.71 |
|  | Levine-BioAge | 60.33 | 59.84 | 4.15 | -0.12 | 9.10E-01 | 1 | 76.39 | 74.81 | 3.19 | -0.49 | 6.20E-01 | 1 | 62.21 | 54.39 | 8.92 |
| Age<br>Acceleration | Horvath-EAA | -14.44 | -1.65 | 2.98 | 4.29 | 6.10E-05 | **1.05E-03** | -2.00 | 0.06 | 1.56 | 1.32 | 1.90E-01 | 1 | 22.74 | 27.95 | 3.67 |
|  | Hannum-EAA | -16.79 | -6.96 | 2.44 | 4.02 | 1.50E-04 | **2.63E-03** | -9.39 | -5.04 | 1.59 | 2.73 | 7.50E-03 | 1.28E-01 | -2.85 | 4.54 | 4.13 |
|  | Zhang.BLUP-EAA | -6.95 | 2.36 | 2.06 | 4.51 | 2.80E-05 | **4.80E-04** | 3.01 | 3.88 | 0.96 | 0.91 | 3.60E-01 | 1 | 3.70 | 7.48 | 1.51 |
|  | Levine-EAA | -15.49 | -7.36 | 3.17 | 2.56 | 1.30E-02 | 2.17E-01 | 1.68 | 3.68 | 1.87 | 1.07 | 2.90E-01 | 1 | 18.27 | 27.04 | 6.85 |
| Intrinsic Age<br>Acceleration | Horvath-IEAA | -14.31 | -4.16 | 2.71 | 3.74 | 3.90E-04 | **6.70E-03** | -2.13 | -1.23 | 1.60 | 0.56 | 5.80E-01 | 1 | 13.13 | 13.27 | 3.91 |
|  | Hannum-IEAA | -9.27 | -1.85 | 2.33 | 3.18 | 2.30E-03 | **3.87E-02** | -2.11 | 1.18 | 1.65 | 2.00 | 4.90E-02 | 8.31E-01 | -4.24 | -1.47 | 3.12 |
|  | Zhang.BLUP-IEAA | -10.35 | -3.92 | 2.98 | 2.16 | 3.40E-02 | 5.84E-01 | 6.65 | 7.90 | 1.92 | 0.66 | 5.10E-01 | 1 | 17.13 | 22.64 | 6.55 |
|  | Levine-IEAA | -7.49 | 0.24 | 2.04 | 3.79 | 3.40E-04 | **5.72E-03** | 2.31 | 2.50 | 1.00 | 0.19 | 8.50E-01 | 1 | -2.64 | -1.87 | 1.69 |
| Extrinsic Age<br>Acceleration | Hannum-EEAA | -7.80 | -0.74 | 2.17 | 3.26 | 1.80E-03 | **3.07E-02** | 3.13 | 6.74 | 1.66 | 2.18 | 3.20E-02 | 5.43E-01 | -2.34 | -0.38 | 2.81 |
|  | Zhang.BLUP-EEAA | -6.17 | -0.91 | 2.79 | 1.89 | 6.30E-02 | 1 | 7.45 | 7.67 | 1.96 | 0.11 | 9.10E-01 | 1 | 20.61 | 24.53 | 5.43 |
|  | Levine-EEAA | -4.80 | 2.53 | 2.04 | 3.60 | 6.20E-04 | **1.06E-02** | 4.94 | 5.01 | 1.01 | 0.07 | 9.50E-01 | 1 | -0.81 | 0.04 | 1.69 |

*Bold numbers identify statistical differences between groups.* *Positive estimate values represent age acceleration values and negative ones indicate age deacceleration.*

### Sex Dichotomy in Age Acceleration in IS Patients

As observed in our previous work ^7^, in this new IS cohort we continue to see how women are chronologically older than men (p-value = 2.3×10^-04^) but have the same biological age (Model.1: Horvath-BioAge, p-value = 1; Hannum-BioAge: p-value = 1; BLUP-BioAge: p-value = 1; Levine-BioAge: p-value = 1) (Supplemental material Table S4). When assessing acceleration measures, we continue to observe lower EAA, IEAA and EEAA values in women than in men, especially for Horvath-EAA, bonf.p-value = 2.6×10^-03^; Hannum-EAA, bonf.p-value = 7.1×10^-03^; Horvath-IEAA, bonf.p-value = 1.6×10^-02^ and Hannum-IEAA, bonf.p-value = 4.5×10^-02^ (Supplemental material Table S4).

### Sex Dichotomy in Age Acceleration control samples

In the specificity analysis, control samples had no differences in chronological age nor any of the biological age measures in the univariate or the multivariate analysis (Supplemental material Table S4). When adjusting for smoking habit (Model.1), as in the ICH analysis, no differences were observed for EAA and when accounting for all potential confounders found in controls (Model.4 – smoking and hypertension) small differences were identified in EAA in which women were slightly deaccelerate compared to men (Horvath-EAA, bonf.p-value = 3.2×10^-02^) and interestingly Hannum-EAA presented, for both sexes, negative acceleration values, although women more so than men (Hannum-EAA, bonf.p-value = 1.8×10^-02^). For both Model.1 and Model.3 analysis, the same patter persisted for IEAA and EEAA measures, in which both sexes had negative acceleration values that were more prominent in women than in men, (Supplemental material Table S4).

As we have seen, women tend to have negative acceleration values regardless of their stroke status. What we observed when comparing EAA from ICH female samples with non-stroke females, is that ICH women are on average more deaccelerated (Horvath-EAA, bonf.p-value = 3×10^114^; Hannum-EAA, bonf.p-value = 4.3×10^-06^; BLUP-EAA, bonf.p-value = 3.6×10^-114^; Levine-EAA, bonf.p-value = 1) behavior that extends to IEAA for biological clocks but only significant for BLUP (BLUP-IEAA, bonf.p-value = 6.9×10^-04^) but is reversed for Levin’s phenotypic clock although not significant (Supplemental material Table S5). Similar results are observed when we compare ICH males with non-stroke male samples, although biological age differences are present (Horvath-BioAge, bonf.p-value = 8.69×10^-03^, Hannum-BioAge, bonf.p-value = 2.26×10^-03^; BLUP-BioAge, bonf.p-value = 4.32×10^-02^; Levine-BioAge, bonf.p-value = 2.8×10^-05^), no statistically significance differences are observed for the acceleration values, with the exception of BLUP-EAA (bonf.p-value = 4.56×10^-04^) (Supplemental material Table S5).

## DISCUSSION

DNA methylation has become a hallmark of aging with a growing body of literature supporting an age specific drift of DNAm patterns[8, 9, 33]. DNAm clocks can precisely calculate someone’s age with a median error of less than four years[26–28], despite that, age acceleration does not seem to be fixed and is depended on several factors. Moreover, contrary to previous belief in which age was thought to be a non-modifiable risk factor, thought modification of DNAm levels biological age can be changed, providing a therapeutic target for disease prevention and management[20].

Here we evaluated three age-predictor clocks. We observed that, despite being chronologically older, women have the same biological age than men at time of ICH onset. Therefore, women have on average lower EAA values than men. The same observation was made for IS patients in a previous study of our group[7], in which women showed negative acceleration values, whereas men had positive ones. These results were replicated in this work, in a new set of patients. This suggests that stroke onset occurs at the same age, biologically speaking, in women and men, but raises the question of what the physiological drivers of such events might be.

More refined metrics of aging, IEAA and EEAA, calculate cell-intrinsic proprieties of aging process independent of changes in blood cell counts, and cell-composition dependent measures of aging, respectively[12]. We observed that sex differences in Horvath-IEAA and Hannum-IEAA, and Hannum-EEAA, mirroring what was observed previously for IS samples[7]. Therefore, differences between women and men might be driven by mechanisms both dependent and independent of cell composition, thus preserve across various cell types and organs, but also capturing aspects of immunosenescence.

It is notable that females with ICH show lower acceleration values compared to those without a stroke. This could be due to a combination of factors, such as an unusual alteration of DNAm-age-related CpG sites during the acute phase of stroke, or because the non-stroke samples are still relatively young, and we may be observing early signs of future brain health issues that have not yet manifested. River et al., using data from 4,000 participants in the Health and Retirement Study[34], found that DNAm age not only reflects past brain health events, such as stroke, dementia, or depression, but also that an elevated biological age significantly increases the risk of these events in the following four years[35].

To ascertain that we were no exclusively identifying pathomechanisms of stroke’s acute phase, we also interrogated Levin’s health-status clock, that was trained as a predictor of mobility and mortality[29]. No significant differences in EAA, IEAA or EEAA were observed between sexes, suggesting that sex differences in acceleration are more specific of the aging process and less related to DNA damage response, increase activation of pro-inflammatory and interferon pathways and decreased activation of transcriptional/translational mechanisms.

A study conducted in over 4.500 samples, uncovered that EEAA had many more associations with a wide range of lifestyle factors, including diet, physical activity, toxic habits and education than IEAA[18]. Consistent with the hypothesis that cell-intrinsic aging remains relatively stable and not influenced by external factors. Another study suggests that women’s cardio protection is lost under conditions of obesity and type 2 diabetes mellitus[36]. It would be interesting to know, in the lifestyle study, the strength of the sex-segregated associations, if women deacceleration is benefited from extrinsic factors compared to men might explain a portion of the differences we are observing.

Another common line of thinking is that sex dimorphism, especially in cardiovascular diseases, are in part driven by female hormonal changes. Especially because the risk of cardiovascular diseases increases in women after menopause[1, 2, 37]. Levie et al. in a three-cohort meta-analysis observed that menopause accelerates epigenetic aging, in blood samples but not in saliva and buccal epithelium samples[38]. The authors suggest that this is because blood samples are better equipped to capture the specific aging process that occurs in women undergoing menopause, and potentially being the reason why there is an increased incidence of strokes in women in the older age rang[38]. Despite that, other studies using mendelian randomization analysis investigate the causal role of women’s reproductive health with small-vessel disease and found contradictory evidence of age at menarche, age at first birth or age at menopause with the risk of ICH or small-vessel IS[39, 40].

When evaluating sex differences by ICH location, we observed that women have lower acceleration values than men in the deep ICH cases. Although, results from the lobar, brainstem and cerebellar subgroups had the same tendency, did not reach statistical significance, most likely due to the smaller simple size. Other studies well powered to perform location-stratified analysis are needed to extract a meaningful conclusion.

Epigenetic modifications are reversible, making them an appealing target for developing therapies aimed at slowing the aging process and reducing the risk of age-related diseases such as intracerebral hemorrhage (ICH). Recent studies have observed that patients treated with common drugs prescribe for age-associated chronic diseases had on average lower EAA measures than those that were not treated[41]. The DIRECT PLUS randomized controlled trial (NCT03020186), interrogated seven DNAm clocks in three interventions, healthy dietary guidelines, a Mediterranean diet, and a polyphenol-rich, low-red/processed meat Mediterranean diet, detecting that both the Mediterranean diet and the polyphenol-rich Mediterranean diet were inversely associated with biological aging[42]. This suggest that targeted therapies could be developed to slow the aging process to mitigating stroke onset as well as personalize therapies to aging women.

There is an extended literature on the sex differences in epidemiological factors in stroke and other disease, but not much is known about the biological age differences. Here we identified differences in biological age acceleration between men and women at time of ICH. Which opens the possibility to developed personalized therapies to slow the aging process based on patient’s sex and other phenotypic characteristics.

## LIMITATIONS

We have evaluated the only ICH cohorts with available DNAm data, which is a rather small cohort to obtain generalized conclusions. Despite that, we’ve been able to observe differences that are in line with what’s described in the literature. It’s important to acknowledge that ICH risk and outcome have an ethnic-dependent component[5, 43], which can also extend to sex differences, thus further assessment of our findings in more diverse cohorts are a must. Most likely, the differences in time-of-blood extraction between the lobar ICH patients of Hospital del Mar and Hospital Sant Pau, has hindered our ability to observe significant differences among groups.

## ACKNOWLEDGMENTS

We thank the Spanish Stroke Genetics Consortium and the RICORS Network.

